# BoxCarmax: a high-selectivity data-independent acquisition mass spectrometry method for the analysis of protein turnover and complex samples

**DOI:** 10.1101/2020.11.20.392043

**Authors:** Barbora Salovska, Wenxue Li, Yi Di, Yansheng Liu

## Abstract

The data-independent acquisition (DIA) performed in the latest high-resolution, high-speed mass spectrometers offers a powerful analytical tool for biological investigations. The DIA mass spectrometry (MS) combined with the isotopic labeling approach holds a particular promise for increasing the multiplexity of DIA-MS analysis, which could assist the relative protein quantification and the proteome-wide turnover profiling. However, the wide isolation windows employed in conventional DIA methods lead to a limited efficiency in identifying and quantifying isotope-labelled peptide pairs. Here, we optimized a high-selectivity DIA-MS named *BoxCarmax* that supports the analysis of complex samples, such as those generated from Stable isotope labeling by amino acids in cell culture (SILAC) and pulse SILAC (pSILAC) experiments. *BoxCarmax* enables multiplexed acquisition at both MS1- and MS2-levels, through the integration of BoxCar and MSX features, as well as a gas-phase separation strategy. We found BoxCarmax modestly increased the identification rate for label-free and labeled samples but significantly improved the quantitative accuracy in SILAC and pSILAC samples. We further applied BoxCarmax in studying the protein degradation regulation during serum starvation stress in cultured cells, revealing valuable biological insights. Our study offered an alternative and accurate approach for the MS analysis of protein turnover and complex samples.

---

Mass spectrometry (MS) based proteomics has matured into a versatile analytical tool in life sciences ^1^. The current state-of-the-art workflows are generally set up on the systems combining high-performance liquid chromatography and tandem mass spectrometry (LC-MS/MS), aiming to achieve both high sensitivity and high selectivity in a reasonable sample throughput. Typically, either high-resolution MS1 profiles of the peptide precursors or high-resolution MS2 profiles of the peptide fragment ions are acquired with sufficient analytical time as bases for peptide quantification ^2,3^.

The MS1-based approach can be exemplified by the accurate mass and retention time tag approach (AMT) ^4^ executed in the latest instruments, in which MS1 spectra of the ultra-high resolution (e.g., 120k or above) are acquired with e.g., an Orbitrap analyzer for the quantitative purpose before MS2-based identification ^5–7^. Meier et al. recently developed an advanced workflow of MS1 profiling named BoxCar ^8^. They decomposed the total peptide ion count across the full MS1 spectrum into narrow m/z segments, each of which was defined by rectangular BoxCar patterns for mass selection. This strategy was built on the ability of the quadrupole-Orbitrap mass analyzer to be filled sequentially with different BoxCar windows that were then analyzed together in a single scan ^8^. The mean ion injection time was reported to increase by >10-fold as compared to that of a standard MS1 scanning, significantly enhancing the MS1-level ion features ^8^. Both MS1 dynamic range and signal-to-noise (S/N) ratios were therefore increased. It should be noted, however, that in BoxCar mode multiple BoxCar MS1 scans are placed in every acquisition cycle, shortening the following MS2 acquisition time.

The MS2-based approach can be exemplified by the data-independent acquisition mass spectrometry (DIA-MS) ^3,9^ executed in the latest instruments. By the guidance of a window table for MS1 isolation covering the full precursor m/z range (rather than the real-time intensity of a peptide precursor ion in a traditional Data dependent acquisition, DDA), DIA-MS essentially acquires the full MS2-level ion features above the detection limit of the mass spectrometer ^3,9^. Recently, DIA-MS has emerged as a popular and powerful method for the reproducible measurement of proteins and proteomes ^1,10,11^. As compared to BoxCar described above, in DIA mode multiple MS2 scans are placed in every acquisition cycle. Each MS2 scan contains mixed fragment ions generated from all peptide ions isolated in a DIA-MS1 window. Due to the limit of scanning speed of the mass spectrometers, the DIA-MS1 window is often wide (e.g., > 10-20 m/z) ^12^.

Because biological and clinical samples frequently generate a complex analytical matrix, the small DIA window settings have been tested in previous literature ^13, 14^. Using smaller windows normally increases the measurement selectivity and reduces the number of co-fragmented precursors in the complex samples ^12, 15, 16^. Nevertheless, using smaller windows sometimes leads to less injection or measurement time per MS2 scan in Orbitrap platforms, which might compromise sensitivity. Egertson et al. proposed an intelligent DIA strategy termed multiplexed MS/MS (MSX), in which five separate 4-m/z isolation windows are analyzed per MS2 spectrum ^17^. By utilizing an additional de-multiplexing step, MSX was reported to significantly increase the DIA-MS selectivity on a level comparable to using 100 4-m/z-wide windows ^17^.

Herein, we argue that the selectivity of DIA method should be given a particular consideration for samples of high-complexity. The samples derived from Stable isotope labeling by amino acids in cell culture (SILAC) ^18^ present such a case. In SILAC analysis, peptide pairs normally ending with heavy (H) and light (L) isotopes are always co-eluted and are, in most cases, measured in the same DIA window^15^. Despite of this difficulty, others and we have previously demonstrated that DIA-MS combined with SILAC ^15,19–21^ and pulse SILAC (pSILAC) experiments ^22–26^ impressively retains the quantitative performance of DIA, as compared to MS 1 −based SILAC approaches ^20, 24^ Recently, we optimized a computational framework to retrieve more H/L features in early time points during pSILAC labeling, for the purpose of quantifying the proteome-wide protein turnover rates ^24^. ***In the present study, we aim to further optimize a particular DIA-MS strategy that supports the analysis of complex samples, such as those generated from SILAC and pSILAC experiments*.** We reason that BoxCar and MSX, in essence, are two methods performing multiplexed data acquisition at MS1 and MS2 levels respectively and can be potentially combined. Thus, we devised a DIA method BoxCarmax, embracing the high-sensitivity of BoxCar ^8^ and the high-selectivity of MSX ^17^. We then demonstrated the utility and superior performance of BoxCarmax in analyzing high-complexity samples and in measuring protein turnover during a cell starvation process.

## MATERIALS AND MERHODS

### Materials and Chemicals

Hela standard peptide digest was purchased from Thermo Fisher (Pierce™, #88328). The human plasma sample was purchased from Sigma (#P9523). Heavy L-Arginine-HCl (13C615N4, purity >98%, #CCN250P1), and L-Lysine-2HCl (13C615N2, purity >98%, #CCN1800P1) were purchased from Cortecnet. RPMI media lacking arginine and lysine was purchased from Thermo Fisher (# 88365).

### Cell culture and pSILAC experiment

Ovarian cancer cell A2780 (#93112519-1VL, Sigma) was cultured for at least eight passages in SILAC media to reach > 99% labeling efficiency (checked by MS). PC12 cells (#CRL-1721, ATCC) were synchronized ^27^ before switching to serum-free SILAC medium. PC12 cells were harvested at 30min, 1h, 4h, 12h, 24h, 48h, and 72h with two dish replicates. Protein extraction and digestion were performed as previously described ^15, 23^.

#### Mass spectrometry measurements

##### LCseparation

For all MS injections, 1 μg of peptide mixture was used. The HPLC separation was performed on EASY-nLC 1200 systems (Thermo Scientific) using an analytical PicoFrit column (New Objective, 75 μm × 50 cm length) self-packed with ReproSil-Pur C18-Q 1.9 μm resin (Dr. Maisch GmbH).

##### 4hr-DIA

The Orbitrap Fusion Lumos Tribrid mass spectrometer (Thermo Scientific) was coupled with a NanoFlex ion source keeping the spray voltage at 2000 V and heating capillary at 275 ° C ^28^. The DIA-MS method for 4hr-DIA consisted of a MS1 survey scan and 33 MS2 scans of variable windows ^29^. The MS1 scan range is 350–1650 m/z and the MS1 resolution is 120 K at m/z 200. The MS1 full scan AGC target value was set to be 2.0E6 and the maximum injection time was 50 ms. The MS2 resolution was set to 30,000 at m/z 200 and normalized HCD collision energy was 28%. The MS2 AGC was set to be 1.0E6 and the maximum injection time was 50 ms.

##### BoxCarmax

In the prototype BoxCarmax presented in this work, the MS1 isolation windows and MS2 scanning m/z ranges are matched in each of the four injections (Injection 1st, 2nd 3rd, and 4^th^, **Figure S1** and **Supplementary Table 1**). The superimposition of the MS1 scans of all four injections reconstructs a full MS1 scan. In each injection, the MS1 scanning is essentially executed through a Targeted Selected Ion Monitoring (t-SIM) scan ^8^ based on quadruple isolation multiplexing. For tSIM MS1, the AGC was set to be 2.0E6 and the maximum injection time was set at 256 ms, and the total AGC target is split equally between the ten different ion species of a multiplexed group. For tSIM, the orbitrap filling is non-isochronous, so that longer injection time is spent for m/z region of low abundance ion species ^8^. The MS2 scanning is executed through tMS2 multiplexing mode (i.e., MSX), in which the MS uses the quadruple to sequentially isolate each targeted ion and fragments them as they enter the ion routing multipole (IRM), simultaneously stores all fragments from the defined number of multiplexed ions (N=4 in each MSX of BoxCarmax) in the IRM, and transports them into Orbitrap for simultaneous mass analysis. The MS2 AGC was set to be 1.0E6 and the maximum injection time was 50 ms. The MS2 resolution was set to be 30,000 and the MS2 scan range was 200-1800 m/z. Please refer to **Figure S2** and further description in the Results.

### DIA data analysis

All DIA analyses including library generation, targeted data extraction, and data reporting were performed based on Spectronaut v14 (both peptide and protein FDR were controlled at 1%) ^30, 31^. The comprehensive sample-specific libraries were generated based on previous publications and datasets in the present study (**Supplementary Experimental Section)**. To perform SILAC and pSILAC targeted data extraction, the Inverted Spike-In workflow was used ^24^. The PECA pS model in the PECAplus Perseus plugin^32^ (https://github.com/PECAplus/Perseus-PluginPECA) was used to estimate the degradation rate parameters from pulsed proteomics data per gene per measurement interval. The hierarchical clustering analysis (HCA), heatmap visualization, gene ontology biological processes (GOBP) annotation, and Fisher’s exact test were performed in Perseus *v1.6.14.0* ^33^. All boxplots and bubble plots were generated using R package “ggplot2”.

### Data availability

The mass spectrometry data and spectral libraries have been all deposited to the ProteomeXchange Consortium via the PRIDE ^34^ PXD021922. (Data will be released upon publication).

See **Supplementary Experimental Section** for details.

## RESULTS AND DISCUSSIONS

### Establishment of BoxCarmax DIA method

For analyzing samples of high complexity, we aimed to develop a DIA-MS method with a high selectivity that is comparable to when windows as small as ~2.5 m/z-wide are used. We modified the previous MSX acquisition ^17^ for this purpose (see below). In the meanwhile, to maintain the detection sensitivity, we aimed to incorporate the enhanced MS1 features offered by the BoxCar concept ^8^. Therefore, this <u>BoxCar</u> and <u>M</u>S<u>X</u> <u>a</u>ligned DIA method was termed as *BoxCarmax* (**Figure 1 and Methods**). BoxCarmax was configured as following: **(1)** The full MS1 spectrum was decomposed into four sets of multiple narrow m/z segments, which were interspaced and subsequent to each other (**Figure 1A**). Each set contained e.g., ten rectangular 22 m/z-wide BoxCar windows. Unlike the original BoxCar method in which all BoxCar window sets were sequentially analyzed in one data acquisition cycle, in BoxCarmax they were respectively analyzed via one of the four injection replicates (i.e., 1st, 2nd 3rd, and 4th injection) of the same sample. **(2)** In each injection, the DIA MS2 measurements were directed to match the corresponding BoxCar MS1 isolation window set. This means, e.g., in the 1st injection, m/z ranges corresponding to BoxCar windows in 2-4th injections are *not* isolated *nor* measured with any MS2 data (**Figure 1B&C**). **(3)** In each of the four injections per sample, the DIA MS2 acquisition was arranged by a total of 30 sequential MSX acquisitions. Four 2.5 m/z isolation windows were multiplexed for each MS2 scan (i.e., each MSX). Contrary to the original MSX method in which the 4 m/z isolation windows were randomly gathered for multiplexing ^17^, in BoxCarmax four 2.5 m/z windows were selected with a fixed, long interspace (i.e., 30×2.5 m/z, 75 m/z) between each other for multiplexing throughout the LC gradient (**Figure 1B&D**). Taken together, by combining the four injections, a total of 480 (i.e., 4 plex × 30 MSX scans × 4 injections) windows of 2.5 m/z were essentially covered by BoxCarmax measurement for one sample (**Figure S2 & Supplementary Table 1**). The MSX strategy used in BoxCarmax cannot distribute ion noises between scans ^17^, but it eliminates the need of the de-multiplexing step and renders other applications (see below) ^17^. In the meanwhile, it is fully compatible to both peptide-centric and spectrum-centric DIA data analysis algorithms ^35, 36^. For example, for a given peptide of interest, its mass spectrometric assay in the spectral library can be retrieved for targeted data extraction in one of the four injections ^3,37^. Both MS 1 and MS2 data extracted from BoxCar and MSX scans can be used for subsequent peptide identification and quantification (**Figure 1E**).

**Figure 1:**
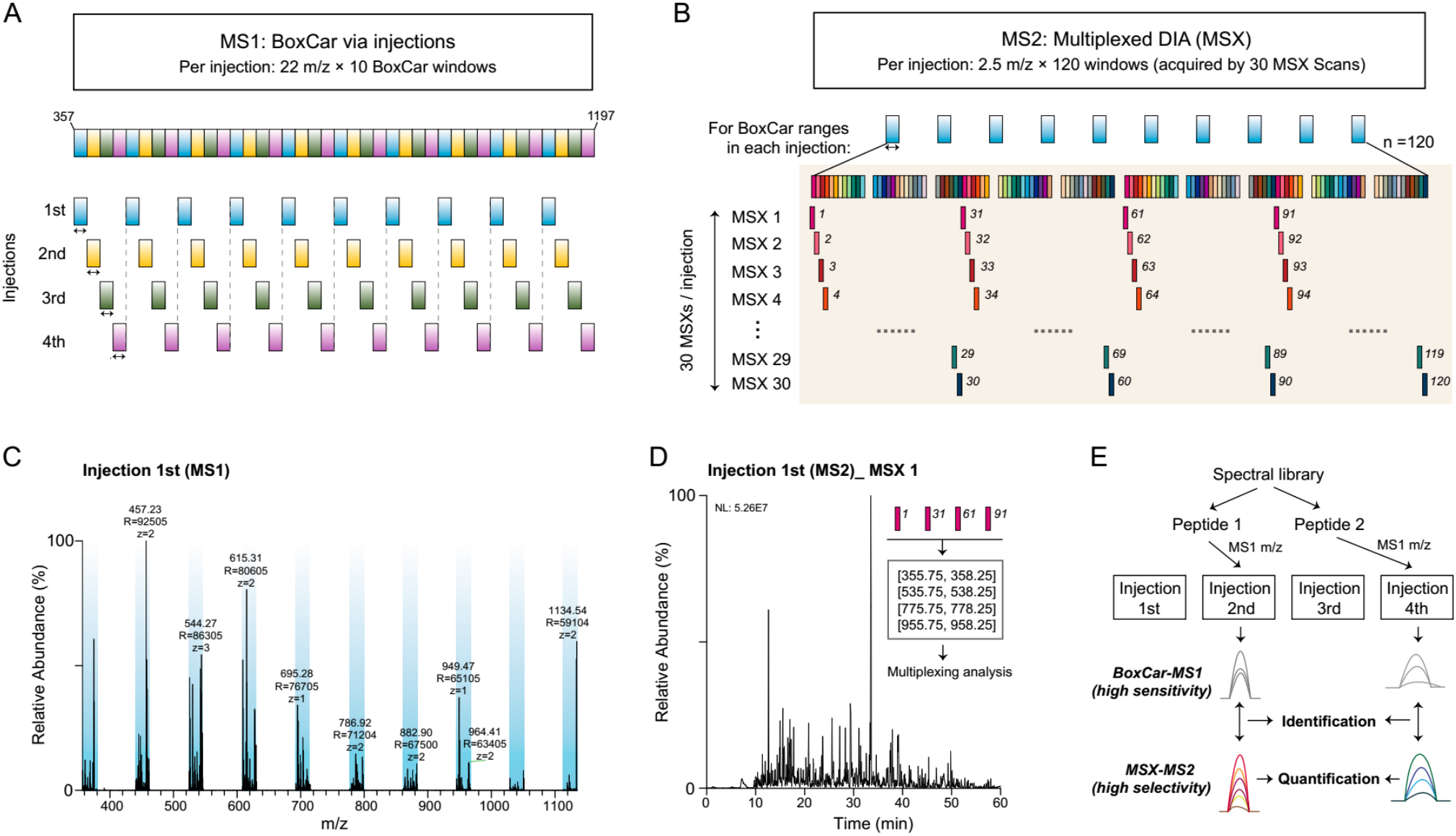
Design of the BoxCarmax workflow for a two-stage multiplexing DIA measurement. **(A)** Schematic representation of MS1 m/z windows in four injections of BoxCarmax. The MS1 scan in each injection has ten rectangular 22 m/z-wide BoxCar windows. The superimposition of the MS1 scans of the four injections reconstructs a full MS1 scan of the sample. **(B)** Schematic representation of the 30 MSX scans in each injection, matching the analytical m/z range of the corresponding BoxCar-MS1 windows. **(C)** A representative example of MS1 scan in Injection 1, with the BoxCar isolation windows shown in blue rectangles. **(D)** Representative example of MS2 signal in the first MSX scan of Injection 1, denoting the four 2.5 m/z-windows multiplexed. **(E)** Example of the peptide-centric identification, which involves MS assay-based data extraction and subsequent peptide identification and quantification.

In summary, we configured BoxCarmax method performing multiplexed analysis at both MS1 and MS2 levels.

### Performance Assessment of BoxCarmax DIA in label-free samples

To assess BoxCarmax performance, we firstly applied it in representative label-free samples, and benchmarked the results by using a state-of-the-art DIA method of 33 variable windows ^29^ that has been actively used in the Liu Lab (**Figure S1**). Considering the practical need of sample throughput, we made each of four BoxCarmax injections one hour and the 33 window DIA a four-hour measurement as the control method (*hereafter*, 4hr-DIA). Thus, a single BoxCarmax analysis requires ~4 hours. The gas-phase separation nature of the four injections indeed identified largely distinctive peptides, with ~85% of total peptide precursors uniquely detected in one of the four injections in a BoxCarmax analysis of a PC12 cell line lysate **(Figure 2A**). The overlapping peptide identifications (~4% of the total between two injections) were due to the design of overlapping m/z edges between adjacent BoxCar MS1 windows (**Supplementary Table 1**). Not surprisingly, many of the distinctively detected peptides in different injections were derived from the same proteins (**Figure S3A**). Because BoxCarmax is essentially a DIA method, as expected, three replicates of BoxCarmax measurement on the same PC12 cell proteome demonstrated excellent reproducibility in both identification and quantification. A total of 84,357 ± 666 unique peptides and 7,927 ± 9 protein groups were identified (FDR = 1% at both peptide and protein levels, same criterial used for all following-up identification results), with a quantitative correlation of R = 0.930-0.953 from the three BoxCarmax replicates (**Figure S3B-D**).

**Figure 2:**
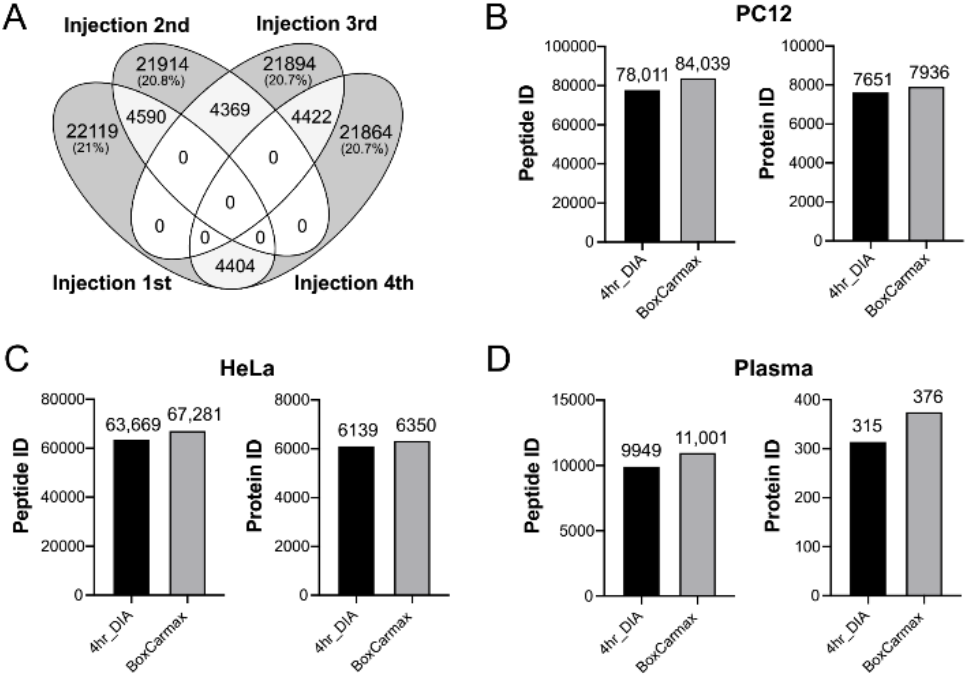
Sensitivity assessment of BoxCarmax in label-free quantification analysis of cellular lysates and plasma samples. **(A)** Venn diagram of peptide precursor ids from Injections 1-4 of BoxCarmax in the PC12 cell line sample. **(B-D)** Number of protein and peptide precursor ids identified by 4hr-DIA method and BoxCarmax in the PC12 cell line lysate **(B)**, HeLa cell line lysate **(C)**, and human plasma **(D)**.

Using respective and extensive sample-specific libraries (see **Supplementary Experimental Section**), we found that BoxCarmax slightly increased the detection rate of 4hr-DIA method. BoxCarmax yielded 7.7% and 3.7% increase of peptide and protein identification numbers in the analysis of PC12 cells, as well as 5.7% and 3.4% increase of peptides and proteins in measuring HeLa standards, as compared to 4hr-DIA (**Figure 2B&C**). Interestingly, when analyzing the human plasma proteome that harbors a much higher dynamic range than cell line or tissue samples ^38^, BoxCarmax identified 10.6% more peptides (11,001 vs. 9,949) and 19.4% more plasma proteins (376 vs. 315) than 4hr-DIA. This indicates the potential of BoxCarmax in analyzing high-dynamic range samples. Taken together, despite of a narrower m/z coverage and 4-times more consumption of peptide samples (**Figure S3**), BoxCarmax retained the favorable quantitative reproducibility of DIA-MS and could serve an alternative DIA method for measuring label-free proteomic samples, especially those of high complexity.

### Advantage of BoxCarmax DIA in analyzing SILAC samples

We reason that the small DIA windows of ~2.5 m/z in BoxCarmax are distant to each other (i.e., ~75 m/z), which are promising for separating the MS2 analysis of a peptide and its variants such as those carrying post-translational modification (PTM) or SILAC labels. Due to the wide isolation windows used in conventional DIA methods, many peptides carrying PTM (such as oxidized methionine) could have been isolated together with the naked peptide target in the same window. This co-isolation may generate DIA peak groups difficult to distinguish, owing to the similar fragmentation patterns between the peptide and its variant forms. Fortunately, the high-performance LC separation and MS1 features (if detectable) could help to distinguish the peaks. However, this issue is especially relevant for SILAC samples in which the isotopic Lysine8 (K8) and Arginine10 (R10) are normally deployed. The H and L peptide counterparts are always co-eluting along the LC gradient. Because K8 and R10 labeling only add small m/z deviations to the light peptides (i.e., 4.007 or 5.004 m/z for charge 2+ precursors, and 2.671 or 3.336 m/z for charge 3+; **Figure 3A**), the H and L peptides are commonly measured by simultaneous MS2 spectrum in most previous DIA approaches whose windows are much larger than e.g., 2.671 m/z, including the original MSX approach ^17^.

**Figure 3:**
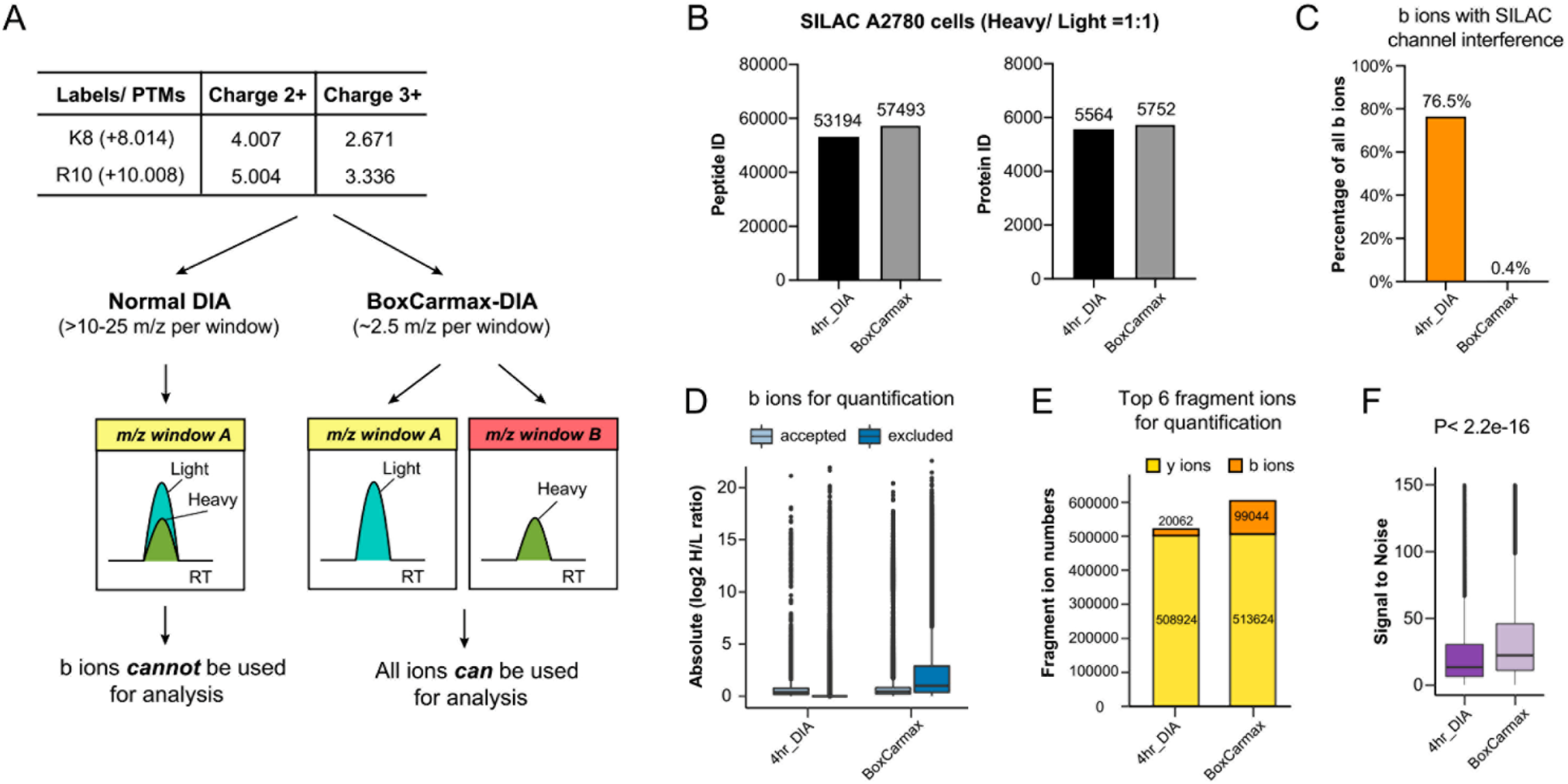
Improved selectivity and b ions usage of BoxCarmax in a SILAC sample of A2780 cells. **(A)** Advantage of BoxCarmax in SILAC analysis, which enables more fragment ions for quantification. **(B)** Number of protein and peptide precursor ids identified by 4hr-DIA and BoxCarmax in the A2780 SILAC 1:1 sample. **(C)** Proportion of b ions with channel interference from all b ions. These ions should not be used for peptide precursor quantification to avoid serious ratio distortion. **(D)** Absolute log_2_ H/L ratio distribution of b ions accepted or excluded from quantification by interference filtering applied by Spectronaut. **(E)** Number of b and y ions retained for quantification after interference filtering in Spectronaut. **(F)** Signal-to-noise distribution of peptide precursors identified by 4hr-DIA and BoxCarmax DIA. The *P*-value was estimated using Wilcoxon test.

To gauge the utility of BoxCarmax in SILAC experiments, we analyzed a sample derived from A2780 cells with a nominal H/L ratio of 1 (based on the spectrophotometer readouts) ^15^. We found that BoxCarmax only identified 8.1% more peptides (57,493 vs. 53,194) and 3.4% more proteins (5,752 vs. 5,564) than 4-hr DIA in this SILAC sample (**Figure 3B**). This represents only a small increase compared to the label-free results, suggesting that the full usage of b-ions could not significantly improve the peptide detection in SILAC-DIA. Conceivably, about 76.4% of all b ions between H and L peptides fell into the same isolation window and were regarded as SILAC channel interference in 4hr-DIA, whereas only 0.4% were regarded as channel interference in BoxCarmax (e.g., charge 4+ peptides) (**Figure 3C**). Despite the limited identification benefit, the overall reduced fragment interference level in BoxCarmax (**Figure S4A**) encouraged us to estimate its quantification performance. The first glance was discouraging, as the 4hr-DIA seemed to generate more peptides with a H/L ratio close to 1 (**Figure S4B**). However, the interference removal function in Spectronaut helped to discern that 60.4% of interfering b ions in 4hr-DIA data had a H:L ratio that was 1:1, which would mislead the interpretation of quantification accuracy if we simply use H/L=1 as a criteria (**Figure 3D,** also see below). In the meanwhile, even after interference removal, BoxCarmax retained 99,044 b ions in Top6 fragment list for the final quantification, which is almost 5 times as many as that in 4hr-DIA (**Figure 3E**). BoxCarmax also offered significantly better signal-to-noise (S/N) scores for peptide identified than 4hr-DIA (**Figure 3F**). To summarize, BoxCarmax was found to slightly improve the analytical sensitivity but significantly reduced the noise and interference levels for quantification in a classic SILAC sample.

### Improved quantification accuracy of BoxCarmax revealed by a pSILAC experiment

To further interrogate the quantitative benefits of BoxCarmax in labeling samples, we performed both BoxCarmax and 4hr-DIA measurements on a cell starvation model labeled by pSILAC. Herein, the serum-free medium was firstly applied to synchronize the PC12 cells (see Methods), a rat pheochromocytoma cell line that was used in enormous pharmacological and signaling transduction studies. The culturing medium was then replaced by a serum-free SILAC (K8R10) version at a time zero (t = 0 hour); and the cell proteome during labeling at the time points of 0.5, 1, 4, 12, 24, 48, and 72 hours was harvested for respective MS measurement. The starvation was therefore included by the serum-free and glucose-consuming medium during this time frame. After 48 hours, a fraction of cells was noticed to detach due to the starvation.

Extrapolating from SILAC result above, we hypothesized that ratio data derived from early pSILAC time points (when H/L ratios were much smaller than 1) could further reveal the quantitative accuracy of BoxCarmax. Reassuringly, such datasets (e.g., t = 4 hours) nicely supported that 4-hr DIA generated more interfering peptide ratios that were close to 1:1, whereas interferences in BoxCarmax did not have such a pattern (**Figure 4A**). Additionally, it should be stressed that the interference removal function in Spectronaut is not available in many other DIA data analysis algorithms – this indicates the flexibility of BoxCarmax for downstream software algorithms. Next, we evaluated the overall accuracy conferred by BoxCarmax in this pSILAC experiment. Due to the residual H/L variations between 0.5-1 hour and the globally disturbed proteome states between 48-72 hours, we focused on the period of 1-24 hours (i.e., four time points) and asked which DIA-MS *generated more peptide ratios that are monotonously increasing* due to the constant pSILAC labeling. Indeed, BoxCarmax quantified more peptides with the correct trend across samples than 4hr-DIA (48,924 vs. 47,772). Furthermore, 85.8% of peptides from BoxCarmax were accepted using the filter of increasing H/L ratio, as compared to 76.7% from 4hr-DIA (**Figure 4B**). Finally, we compared the final quantitative reports *after* interference removal: Intriguingly, BoxCarmax data at all the time points unanimously resulted in lower H/L ratios than that of 4hr-DIA (P < 2.2e-16 for every time point; **Figure 4C**). The ratio difference is even more remarkable at early labeling time points. Therefore, our data demonstrated that BoxCarmax successfully and significantly reduced the quantification noises that would otherwise tend to make the H/L ratio closer to 1.

**Figure 4:**
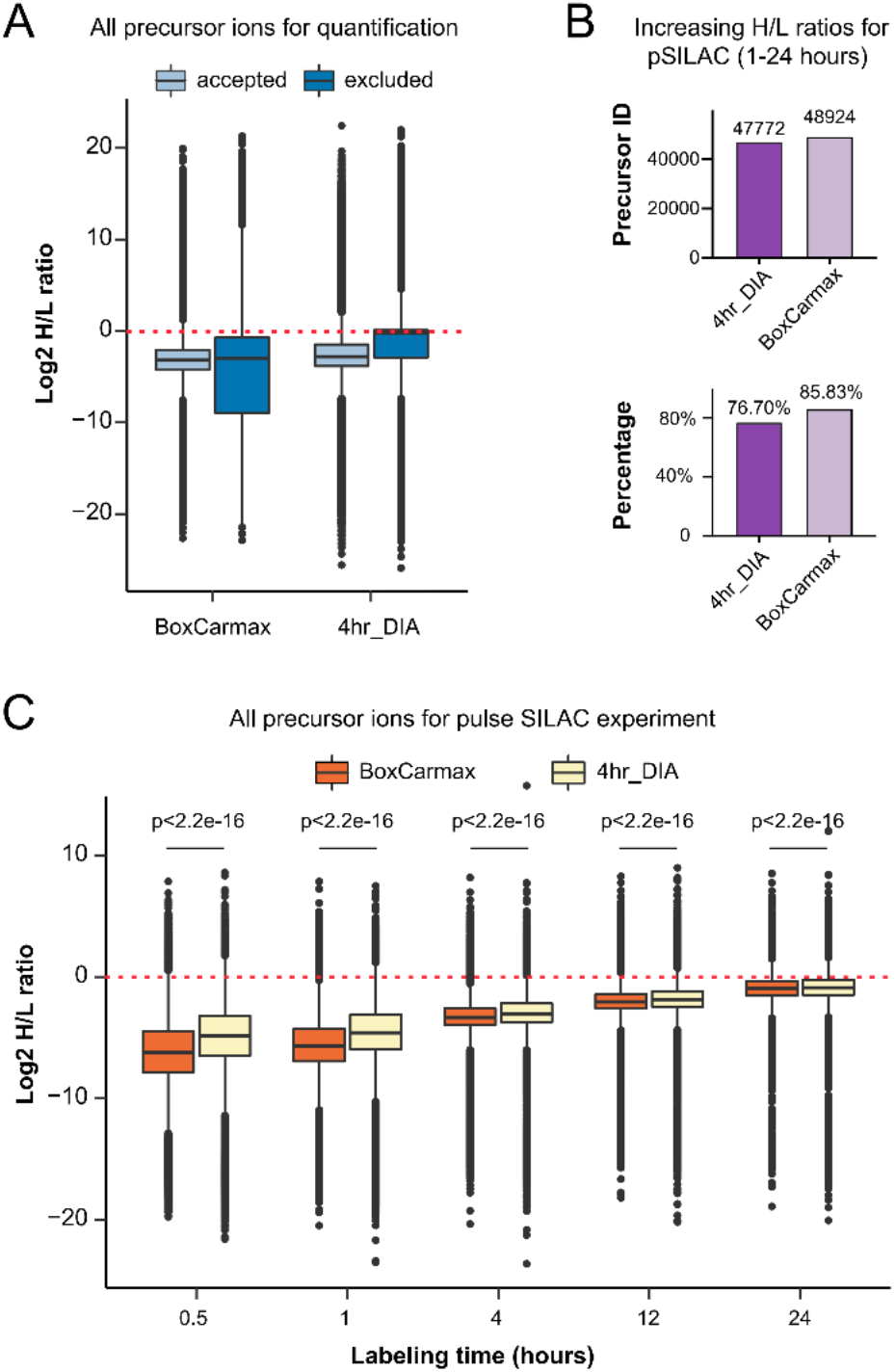
Improved quantification accuracy of BoxCarmax in a pSILAC measurement on PC12 cells. **(A)** Log_2_ H/L ratio distribution of all ions accepted or excluded from quantification by interference filtering applied by Spectronaut in the t = 4 hr time point. **(B)** The number and proportion of peptide precursor ids for which the H/L ratios were monotonously increasing over the 1 hour to 24 hours labeling time points. **(C)** Comparison of precursor H/L ratios in all time points between 4hr-DIA and BoxCarmax DIA. *P*-values were estimated using Wilcoxon test.

### BoxCarmax delineates protein degradation events during cell starvation

Mammalian cells are equipped with elaborate adapting mechanisms for their survival in nutrient starvation ^39–41^. Among different starvation manners, serum starvation is widely deployed in hundreds of biological and pharmacological research studies using culturing tissue cell lines. However, the impact of serum starvation on cellular proteome and proteostasis remains unclear ^42, 43^. The major cellular degradative mechanisms during nutritional deprivation present one of the most fundamental questions in cell biology ^39–41, 44^.

**Figure 4C** illustrated that BoxCarmax improved the overall quantification accuracy in pSILAC experiment especially on early time points, which are essential for robust and accurate protein turnover calculation ^24, 26^. Therefore, we applied BoxCarmax to measure the rate of protein degradation change during the serum starvation time intervals in PC12 cells labeled by pSILAC. *First,* using Protein expression control analysis (PECA) ^33^ on BoxCarmax data, we found that the absolute rates of protein degradation show a global reduction following serum starvation (**Figure 5A**; **Supplementary Table 2**). This likely reflected that, despite the net protein degradation during starvation via ubiquitin-proteasome and autophagy-lysosome pathways ^45^, the cells tried to maintain the cellular proteome hemostasis by attenuating the global speed of degradation (i.e., degradation rate). This mechanism might help the cells to reduce the general cost of protein-level control ^11, 46^, and ensures basic cellular functions in starvation stress. *Second,* we found that individual protein degradation rates did not follow a uniform down-regulation. Instead, clustering analysis based on degradation rates of 3,764 proteins between time intervals (**Figure 5B**) revealed protein degradation was largely variable and could be time- and GO biological process-dependent (**Supplementary Table 3**). This result indicates a delicate dynamic coordination of protein turnover based on their protein function, which is essential to meet the constantly changing demand of the cell. *Third,* since previous literature highlighted that the cellular proteome, when under stress, showed organelle-ordered ^47^ or compartment-specific ^22^ degradation pattern, we herein compared the protein degradation speed during 1-4 hours (D1) and 24-48 hours (D4) between major cellular organelles for a total of 5065 proteins (**Figure 5C**). Whereas many organelles did not show a broad accelerating or decelerating protein degradation relatively, we found a notable faster degradation for mitochondrion (1D enrichment P value= 2.1e-6) ^48^ and ribosome proteins (P = 0.00028), as well as an annealing degradation for centrosome (P = 0.00035) and plasma membrane proteins (P = 3.2e-05), demonstrating that serumstarvation could exert organelle-specific impact on protein turnover. The finding of preferably degraded ribosome during nutrient-stress is consistent to previous reports ^39, 40^. Altogether, the BoxCarmax results uncovered both global and specific protein degradation changes following serum starvation.

**Figure 5:**
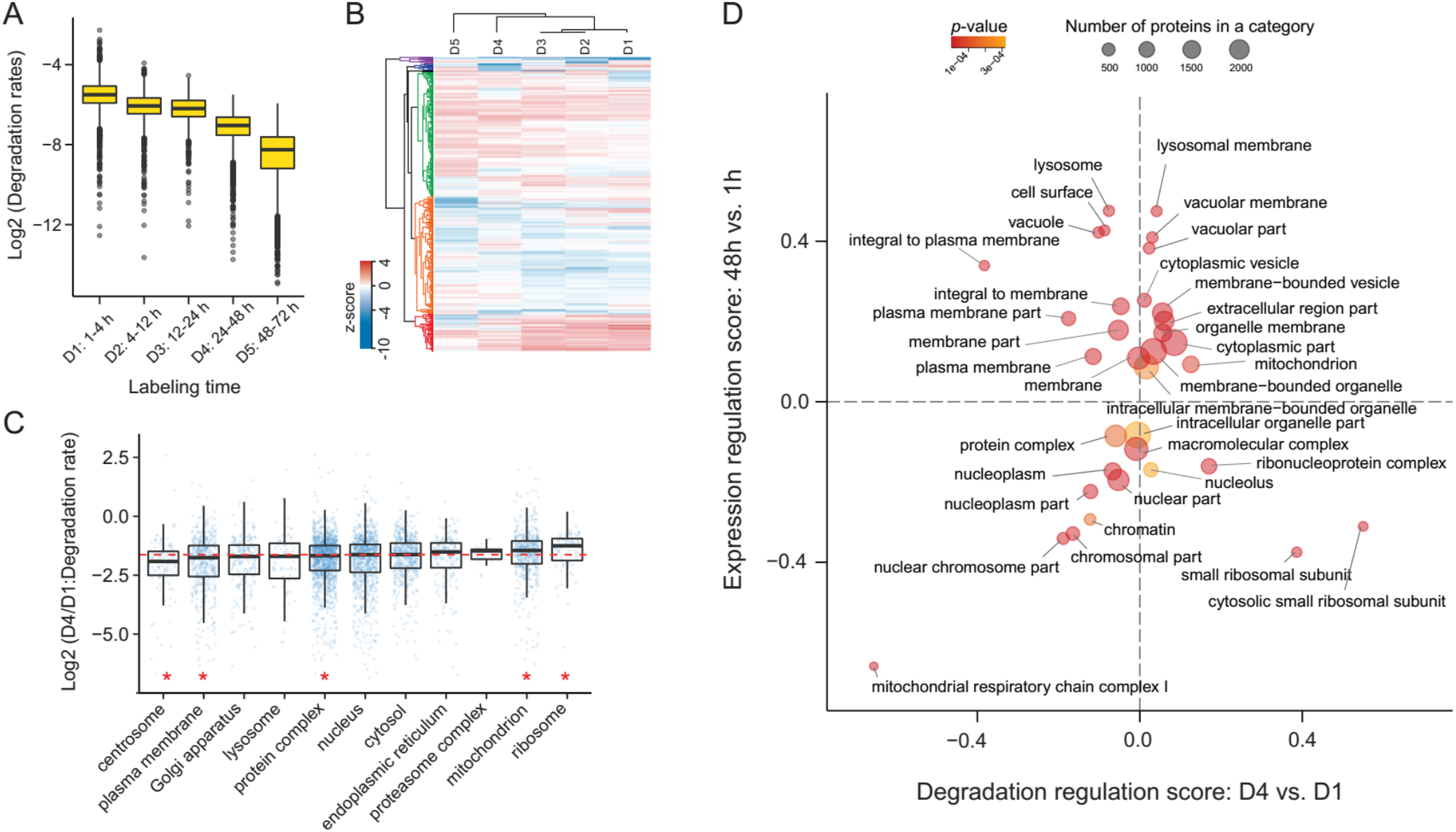
Application of BoxCarmax and pSILAC to study protein degradation in PC12 cells during serum starvation. **(A)** Distribution of log2 degradation rates per indicated time intervals over the time course of the pSILAC experiment in serum-starved PC12 cells. Note the cells were synchronized for 16 hours prior to the heavy isotopic labeling. **(B)** Hierarchical clustering analysis of z-score normalized rates per time interval. D1 to D5 denote the degradation rates between the time intervals. **(C)** Distribution of log2 D4/D1 rate ratios for selected GOCC compartments. Red asterisks indicate statistical significance in 1D enrichment analysis (Benjamini-Hochberg FDR < 0.05). **(D)** Two-dimensional enrichment analysis bubble chart of selected examples of cellular compartments. X-axis represents the enrichment score of log2 D4/D1 degradation rate ratios; Y-axis represents the enrichment score of log2 protein expression fold changes between 48h and 1h. All presented examples were significant in a 2D enrichment analysis (Benjamini-Hochberg FDR < 0.01).

Finally, to inspect whether protein degradation rate measurement brings more biological insights, we added the H and L intensities for each protein, normalized the sum at each time point to represent the total protein abundance, and compared protein abundance change with the degradation change using a 2D enrichment plot ^48^ (Enrichment FDR <0.01, **Figure 5D**, and refer to **Figure S5** for a result with a loosen FDR cutoff of 0.05). This analysis suggested that lysosome proteins upregulated their abundances following starvation significantly as previously reported ^45^, whereas their protein degradation rates were regulated only modestly. In comparison, the down-regulation of ribosome protein abundances was accompanied with a drastic up-regulation of protein degradation during starvation. This diversity emphasizes the different extent of protein translation and degradation contributing to the conventional abundance-centric analysis. Last but not the least, we found certain organelle sub-structures ^24, 44^ could be regulated selectively or distinctively by protein turnover and post-transcriptional process (**Figure S6**). For example, although mitochondrial proteome had a relatively faster protein degradation in the starving cells, the respiratory chain complex I exceptionally reduced protein degradation process, suggesting the post-translational buffering role of protein degradation for complex I to cope with starvation. Thus, the protein degradation profiling provides additional and valuable biological implications that are not evident by analyzing protein abundance alone.

## CONCLUSION

In this work, we developed a particular DIA-MS approach, BoxCarmax, that is highly multiplexing at both MS1 and MS2 levels. BoxCarmax incorporates the sensitivity improvement of BoxCar MS1 acquisition ^8^ through four sample injections and the selectivity improvement of MS2 through non-consecutive acquisition small DIA windows. Similar to a very recent DIA technique based on extensive gasphase separation ^49^, BoxCarmax reaches the sensitivity of nominal isolation width of 2-2.5 m/z, but with a measurement of only four hours per sample due to its multiplexity capacity. Compared to the high-performance classic DIA method (4hr-DIA in this report as an example), BoxCarmax modestly increased the identification rate for label-free and labeled samples but significantly reduced the interference and improved the quantitative accuracy in SILAC and pSILAC samples. Combing with pSILAC, BoxCarmax data deciphered how cells maintain the critical balance between protein synthesis and degradation, to meet the serum depletion stress during tissue culture. This application example thus points out the importance of having the additional control of serum starvation in cell signaling and pharmacology studies.

The limitations of BoxCarmax include the consumption of four-times more peptide samples, which normally does not present a problem in pSILAC experiment using cultured cells. The data points per peak of BoxCarmax in this study are similar to that in conventional 1-hour DIA, which seem to be sufficient for quantification ^50^. Future developments of BoxCarmax-type method may include e.g., the development of non-isochronous filling for both MS1 and MS2 multiplexing via software such as Maxquant.live ^51^ (currently, MSX uses isochronous filling at MS2 level), as well as the development of the corresponding scan-wise correction algorithm enabling relative quantification (if non-isochronous MS2 filling is used). Nevertheless, due to the 2.5 m/z −level selectivity, we expect potential applications of BoxCarmax in analyzing small protein PTMs such as oxidation and methylation, as well as samples of high-dynamic range (e.g., plasma proteomics), high complexity (e.g., metaproteomics), and samples containing lots of peptide isomers (e.g., histone modification analysis).

## AUTHOR INFORMATION

### Notes

**The authors declare no competing financial interest.**

## Acknowledgements

We thank Anatoly Kiyatkin and Archer Hamidzadeh for advices on culturing PC12 cells. We thank Lukas Reiter for technical support in Spectronaut software and George Rosenberger and Hannes Röst for helpful discussions. Y.L. thanks the support from the National Institute of General Medical Sciences (NIGMS), National Institutes of Health (NIH) through Grant R01GM137031 to Y.L.

## Author Contributions

B.S. and W.L. contributed equally to this study. B.S. analyzed the MS data and performed bioinformatics analysis. B.S. prepared most illustrations for the figures. W.L. cultured the PC12 cells and performed all the MS measurements. Y.D. provided critical comments to the manuscript. Y.L. and B.S wrote the paper. Y.L. Y.L conceptualized the method and the study. Y.L. supervised the study.

